# Contamination of Engram Coactivity Networks During Forgetting

**DOI:** 10.64898/2026.04.21.719933

**Authors:** Lulu An, Miyoung Yang, Hongbing Wang

## Abstract

Forgetting is a fundamental component of adaptive memory and essential for cognitive flexibility, yet its cellular basis remains unclear. Here we establish a mouse model of retroactive interference (RI) and show that post-learning novelty exploration induces active forgetting of hippocampus-dependent object location memory only within a discrete consolidation window defined by protein synthesis sensitivity. RI imposed within this window reduces engram reactivation and destabilizes a structured coactivity network formed during learning. Mice that forget retrieve reorganized engram with increased edge turnover and reduced training edge survival during recall. During this vulnerable period, RI infiltrates the engram core, whereas after consolidation it remains confined to the network periphery. Over the consolidation window, the engram network progressively matures, acquiring greater density, similarity, and k-core robustness, features that confer resistance to interference. Importantly, blocking RI infiltration rescues memory formation. Together, these findings show that forgetting arises from reorganization of engram topology during consolidation and identify engram core contamination as a network-level substrate for forgetting.

## INTRODUCTION

Forgetting is an essential activity-dependent process that underpins the efficiency of the learning and memory system (*1-3*). In a dynamic environment where not all information is equally relevant, the ability to prioritize and selectively forget less salient or outdated experiences is vital for optimizing memory and ensuring cognitive flexibility (*4*). This selective forgetting allows for better adaptation and focused cognition, whereas an inability to forget can impair daily functioning. For instance, individuals with highly superior autobiographical memory often struggle to manage routine demands (*5, 6*), and deficits in cognitive flexibility are prominent features of neurological and psychiatric disorders, including traumatic brain injury (TBI), frontotemporal dementia (*7*), post-traumatic stress disorder (PTSD) (*8, 9*), and obsessive-compulsive disorder (OCD) (*10*). Despite its behavioral importance, the cellular and circuit-level mechanisms underlying activity-dependent forgetting remain poorly defined.

Human cognitive studies have proposed retroactive interference (RI), a post-learning activity, as a major driver of memory decay (*3*). In a canonical paradigm, participants first learn a word list as a “to-be-remembered” task and subsequently learn a second word list as the RI task. Data from multiple studies demonstrate that RI induces memory loss (*3*). While these cognitive studies illuminate the psychological dimensions of forgetting, the underlying neurobiology has not been examined.

In this study, we developed behavioral and cellular paradigms to study activity-dependent forgetting in mice and determined the impact of RI on the behavioral and cellular outcomes of memory formation.

## RESULTS

### Temporally constrained RI causes forgetting

We trained mice on a hippocampus-dependent task to learn the locations of two objects. Thirty minutes (0.5 hr) after training, mice underwent 10- or 30-minute novelty exploration (NE) in a novel chamber filled with various objects (e.g., LEGO blocks and small toy figures). Object location memory (OLM) was tested 24 hours later (**Fig. 1A1**). Compared to controls, which showed intact OLM and consequently spent more time exploring the object at the novel location, mice that experienced 30-minute, but not 10-minute, post-learning NE lost OLM (**Fig. 1A2–4**). In contrast, a pre-learning 30-minute NE (**Fig. 1B1**) did not affect OLM (**Fig. 1B2–4**). These data demonstrate that there is a threshold duration for effective interference. Retroactive interference (RI), but not proactive interference (PI), leads to active forgetting.

**Fig. 1.**
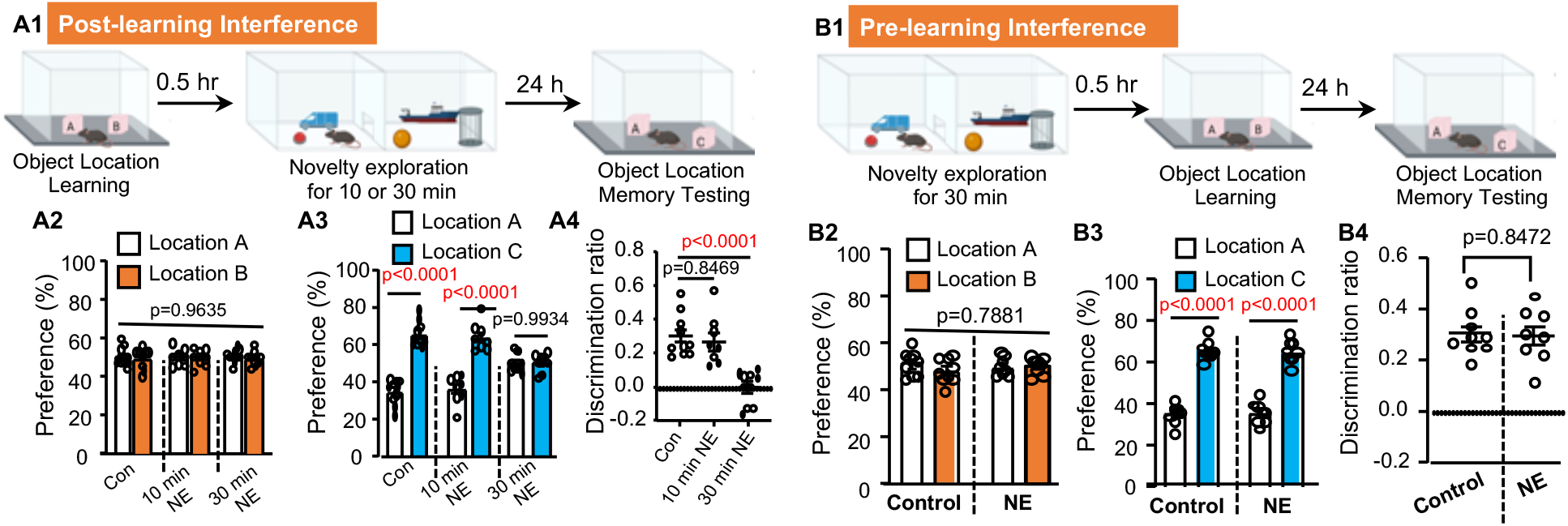
Post-learning novelty exploration causes forgetting of object location memory (OLM). (**A)**. Post-learning exposure to novelty exploration (NE) disrupts OLM (n=9 for the control group; n=8 for the 10 min NE group; n=10 for the 30 min NE group). Mice were trained to learn 2 objects at locations A and B. 0.5 hr after training, mice were exposed to a novel chamber filled with various objects for 10 or 30 min. Mice were tested for OLM 24 hr later (**A1**). During learning, all groups of mice show comparable preference to interact with objects in location A and B [F (2,25)=0.03721, p=0.9635] (**A2**). During testing, the control group (with no novelty exploration) and 10 min NE group but not the 30 min NE group showed preference for the object at the new location C [F (2,25)=18.01, p<0.0001 in (**A3)] [**F (2,25)=17.88, p<0.0001 in (**A4)**]. (**B)**. Pre-learning exposure to NE show no effects on OLM (n=9 for the control group; n=8 for the pre-learning NE group). (**B1)**. Behavior paradigm. (**B2)**. Object preference during learning [F (1,15)=0.395, p=0.5387]. (**B3)**. Object preference during testing [NE effect: F (1, 15) =0.05, p=0.827]. (**B4)**. Object discrimination during testing. Data are presented as mean +/-SEM. Data were analyzed by unpaired t-test (**B4**) and two-way ANOVA (**A2, A3, B2, B3**) or one-way-ANOVA (**A4**) followed by post-hoc pairwise comparison.

Memory consolidation requires new protein synthesis (*11, 12*). We administered the protein synthesis inhibitor anisomycin (150 mg/kg) either 1 or 4 hr after object location learning (**Fig. 2A**). Mice receiving anisomycin at 1 hr, but not 4 hr post-learning failed to demonstrate OLM (**Fig. 2B1–3**), indicating that consolidation is complete within 4 hours of acquisition. Notably, mice exposed to post-learning NE at 0, 0.5, 1, or 2 hr, but not at 4 hr, after training failed to show OLM (**Fig. 3**). Collectively, these data delineate the temporal dynamics of RI-mediated forgetting and implicate a functional interaction between RI and the consolidation process.

**Fig. 2.**
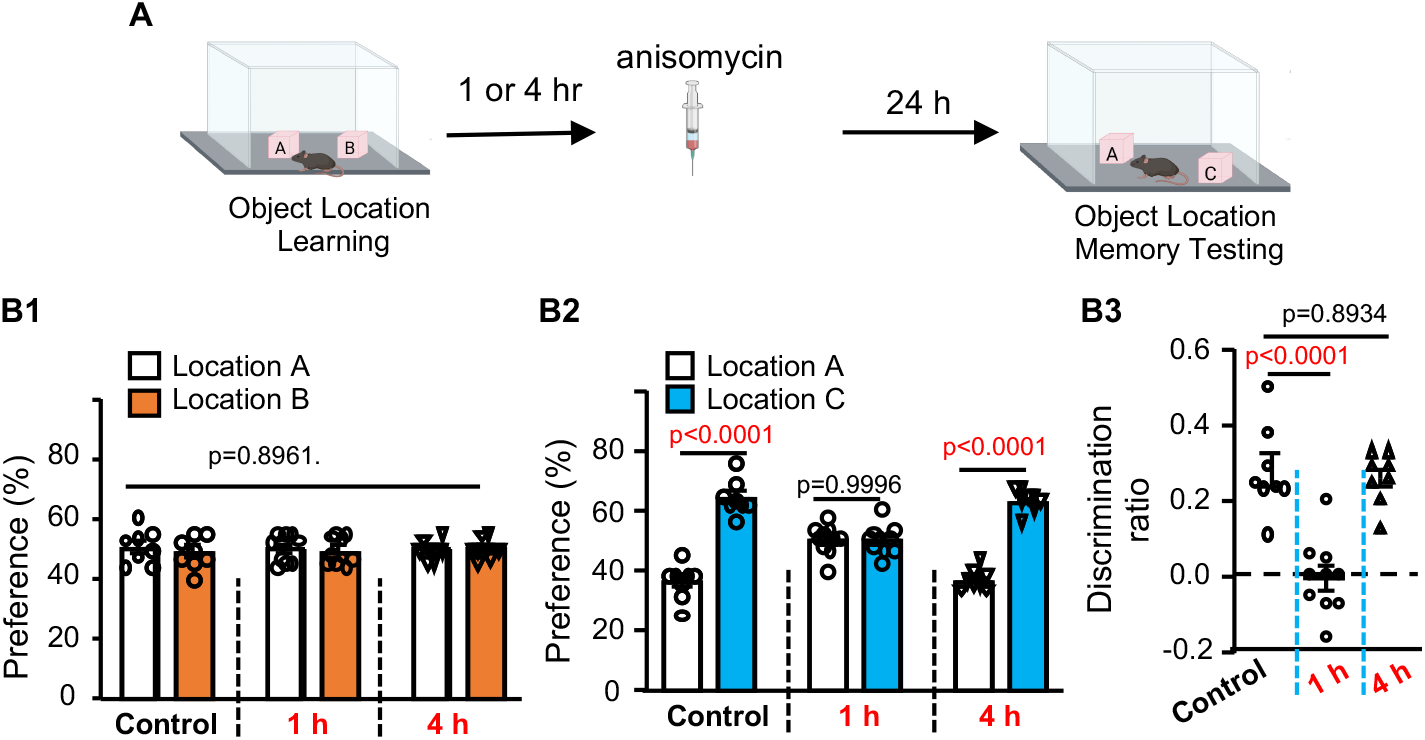
Anisomycin disrupts OLM in a time-dependent manner. **(A)**. Mice were trained to learn 2 objects at locations A and B. The trained mice were then i.p. injected with anisomycin 1 or 4 hr after training (n=8, 9, and 8 for the control, 1-hr, and 4-hr anisomycin groups, repectively). Mice were tested for OLM 24 hr later. All groups showed normal and non-discriminative preference to object at location A and B during training [F (2,23) = 0.1102, p=0.8961] (**B1**). During OLM testing, the control and mice injected with anisomycin at 4 but not 1 hr post-training showed preference to object at the new location C [F (2,23) = 25.24, p<0.0001] (**B2**) and higher discrimination ratio [F (2,23) = 25.15, p< 0.0001] (**B3**). Data are presented as mean +/-SEM and analyzed by two-way (**B1** and **B2**) or one-way ANOVA (**B3**) followed by post-hoc pairwise comparison.

**Fig. 3.**
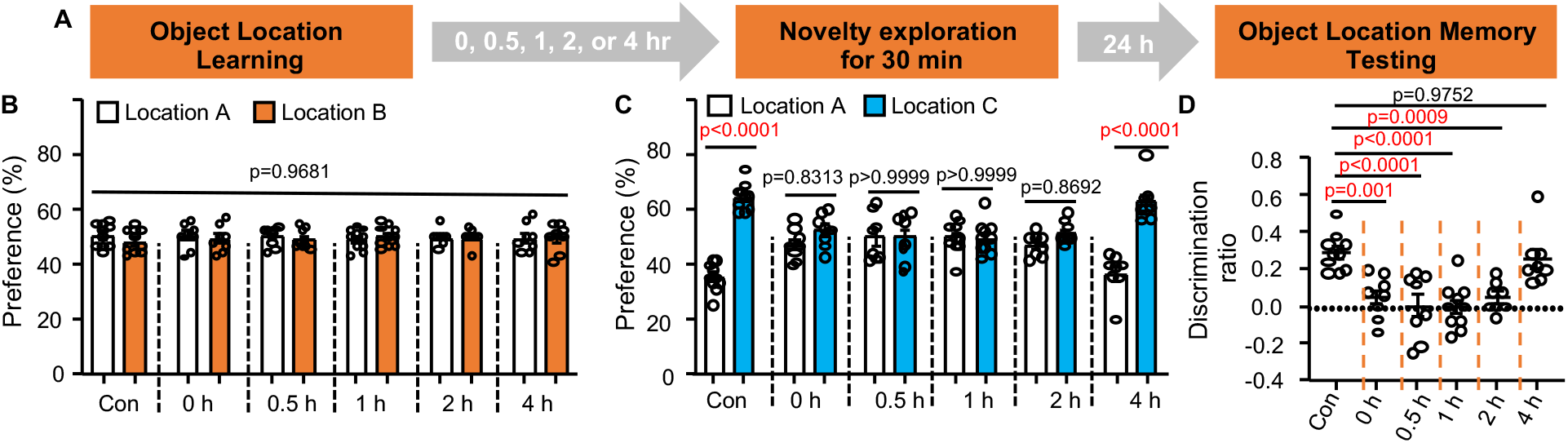
RI within 4 hr after learning disrupts OLM. (**A)**. Behavior paradigm (n=10 for the control and 1-hr NE group; n=8 for other groups). (**B)**. Object preference during learning [F (5, 46)=0.1816, p= 0.9681]. (**C)**. Object preference during testing [F (5,46)=9.999, p<0.0001]. (**D)**. Object discrimination during testing [F (5,46)=9.944, p<0.0001]. SEM. Data were analyzed by two-way ANOVA (**B, C)** or one-way-ANOVA (**D**) followed by post-hoc pairwise comparison.

### Task-specific ensemble during learning, RI, and memory retrieval

To elucidate the cellular mechanisms underlying RI-induced forgetting, we recorded neuronal activity in the dHip using *in vivo* calcium imaging (**Fig. 4A**). An AAV vector was used to drive expression of the calcium sensor jGCaMP7f in dCA1 (**Fig. 4A1**), and a head-mounted fluorescent microscope captured calcium transients and event rates in activated neurons (**Fig. 4A2–3**). We examined three groups of animals experiencing object location learning (training), post-learning activity in home cage (the control group), exposure to NE at 0.5 hr post-learning (the 0.5-hr NE group), exposure to NE at 4 hr post-learning (the 4-hr NE group), and OLM testing (**Fig. 4B1**). Neuronal activity was sampled during training, the post-learning period, novelty exploration, and OLM testing (**Fig. 4B2**). jGCaMP7f expression and the recording procedure did not affect behavioral outcomes. All mice displayed normal learning (**Fig. 4C1**). OLM was intact in the control and 4-hr NE groups but was disrupted in the 0.5-hr NE group (**Fig. 4C2–3**).

**Fig. 4.**
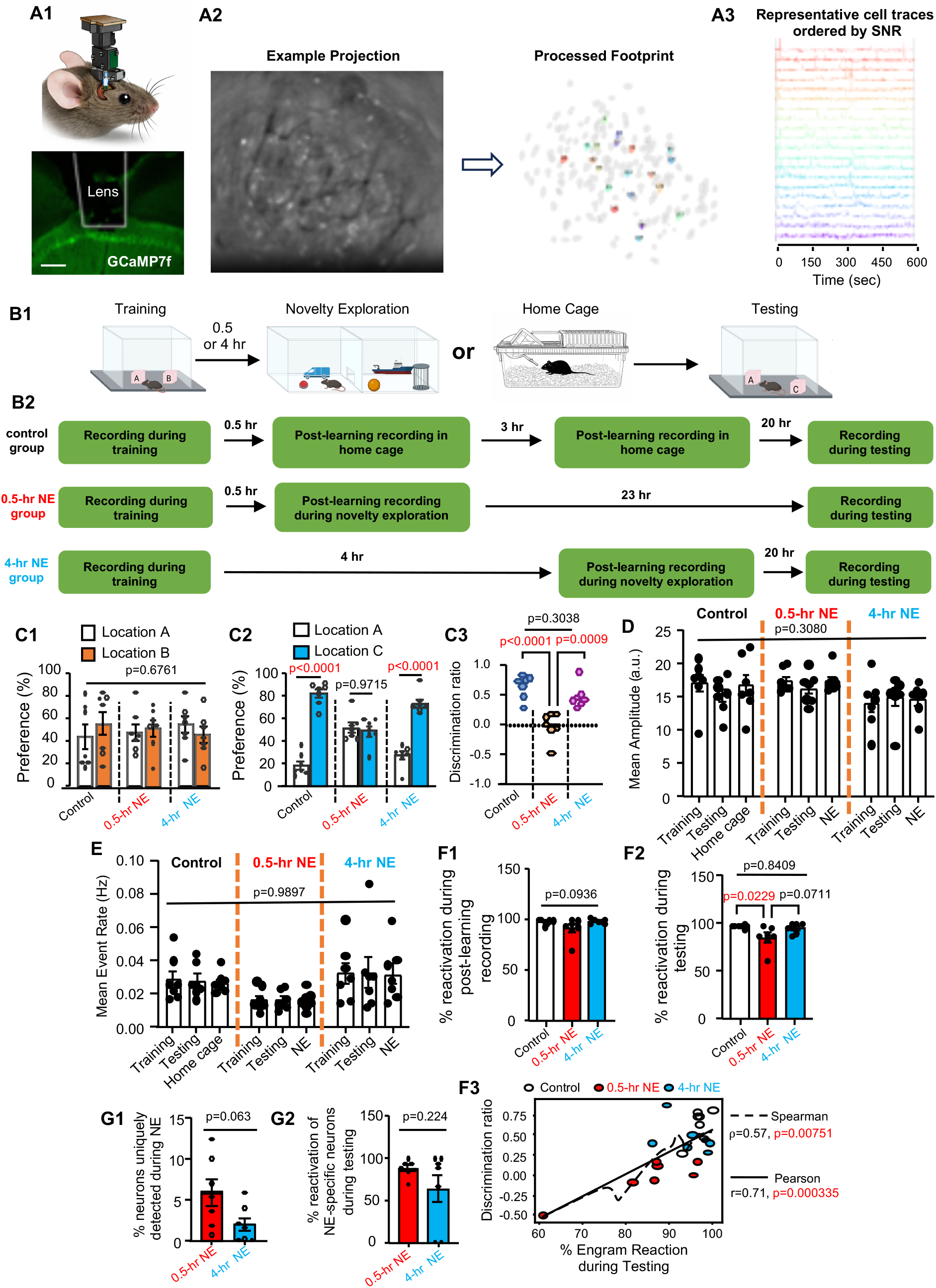
Dynamic activation and reactivation of task-specific ensemble during forgetting. *In vivo* calcium imaging was used to record longitudinal neuronal activity in dorsal CA1 during learning, post-learning NE, and OLM testing. (**A1)**. Schematic of the miniscope placement, lens track, and GCaMP7f expression. Scale bar: 100 µm. (**A2)**. Generation of the processed footprint from raw image by CNMF-E. (**A3)**. Representative cell traces. (**B)**. Behavior procedures (**B1**) along with task-specific calcium imaging (**B2**) with control mice, mice subjected to NE 0.5 hr post-learning (the 0.5-hr NE group), and mice subjected to NE at 4 hr post-learning (the 4-hr NE group). (**C)**. The *in vivo* calcium imaging procedure does not affect learning [**C1**, F (2,18)=0.4, p=0.6761] and the outcome of OLM [**C2**, F (2,18)=19.20, p <0.0001] [**C3**, F (2,18)=19.20, p<0.0001]. While the control and 4-hr NE group showed normal OLM, the 0.5-hr NE group forgot. (**D)**. Mean amplitude during training (all groups), post-training in home cage (control group), post-training NE (0.5-hr and 4-hr NE groups), and testing (all groups) [F (4,36)=1.249, p=0.3080]. (**E)**. Mean event rate during training (all groups), post-training in home cage (control group), post-training NE (0.5-hr and 4-hr NE groups), and testing (all groups) [F (4,36)=0.07365, p=0.9897]. (**F1)**. Ensemble reactivation during post-learning recording [F (2,18)=2.710, p=0.0936]. (**F2)**. Ensemble reactivation during testing [F (2,18)=4.852, p=0.0206]. (**F3)**. Spearman and Pearson analysis were used to determine correlation between discrimination ratio and % of the learning-associated ensemble reactivation during testing. (**G1)**. % of total neurons uniquely detected during NE. (**G2)**. % of reactivation of NE-specific neurons during testing. Data are presented as mean +/-SEM (n=7 for each group). Data were analyzed by unpaired t-test (**G1** and **G2**) and two-way ANOVA (**C1, C2, D**, and **E**) or one-way-ANOVA (**C3, F1**, and **F2**) followed by post-hoc pairwise comparison.

Activated neurons showed comparable calcium transient amplitudes (**Fig. 4D**) and event rates (**Fig. 4E**) across training, home-cage offline periods or NE, and testing. The ensemble manifested during learning was reactivated after training (**Fig. 4F1**) and during OLM testing (**Fig. 4F2**). NE did not affect post-learning ensemble reactivation (**Fig. 4F1**); however, ensemble reactivation during memory retrieval was significantly reduced in the 0.5-hr NE group but not in the control or 4-hr NE groups (**Fig. 4F2**). Spearman and Pearson analyses further identified positive monotonic and linear relationships, respectively, between the degree of ensemble reactivation and memory strength (**Fig. 4F3**).

Calcium imaging also identified NE-specific ensemble, a population activated during NE that was distinct from the learning-activated ensemble (**Fig. 4G1**). These NE-specific neurons were reactivated alongside the learning-associated ensemble during testing (**Fig. 4G2**). Immunostaining with cFos also detected neuron activation in the hippocampus during NE (**Fig. 5**). Intriguingly, although NE-specific neurons in both the 0.5-hr and 4-hr NE groups were reactivated during testing, only the 0.5-hr NE group exhibited an overall reduction in learning-associated ensemble reactivation (**Figs. 4F2–3**) and impaired memory retrieval (**Figs. 3C and 4C**).

**Fig. 5.**
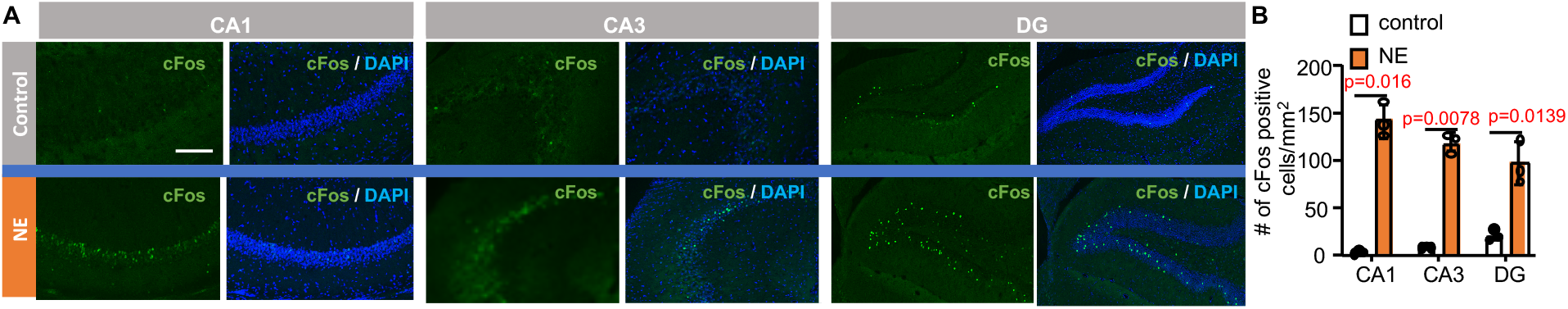
NE causes neuronal activation in the dorsal hippocampus. (**A)**. Hippocampal sections obtained from control and NE mice were immunostained with antibodies against cFos. (**B)**. Quantification of cFos-positive cells in CA1, CA3, and DG of the hippocampus (n=3 for each group). Data are presented as mean +/-SEM. Data were analyzed by unpaired t-test

### RI affects survival and turnover of the engram network

Based on Hebbian principles (*13*), we sought to characterize the memory trace beyond ensemble activity by constructing neuronal coactivity networks (*14, 15*) from learning and retrieval sessions. Coactivity networks capture pairwise neuronal interactions based on coincident calcium events (**Fig. 6A**). We used the training-session coactivity network as a reference and examined network reactivation during testing (**Fig. 6B**).

**Fig. 6.**
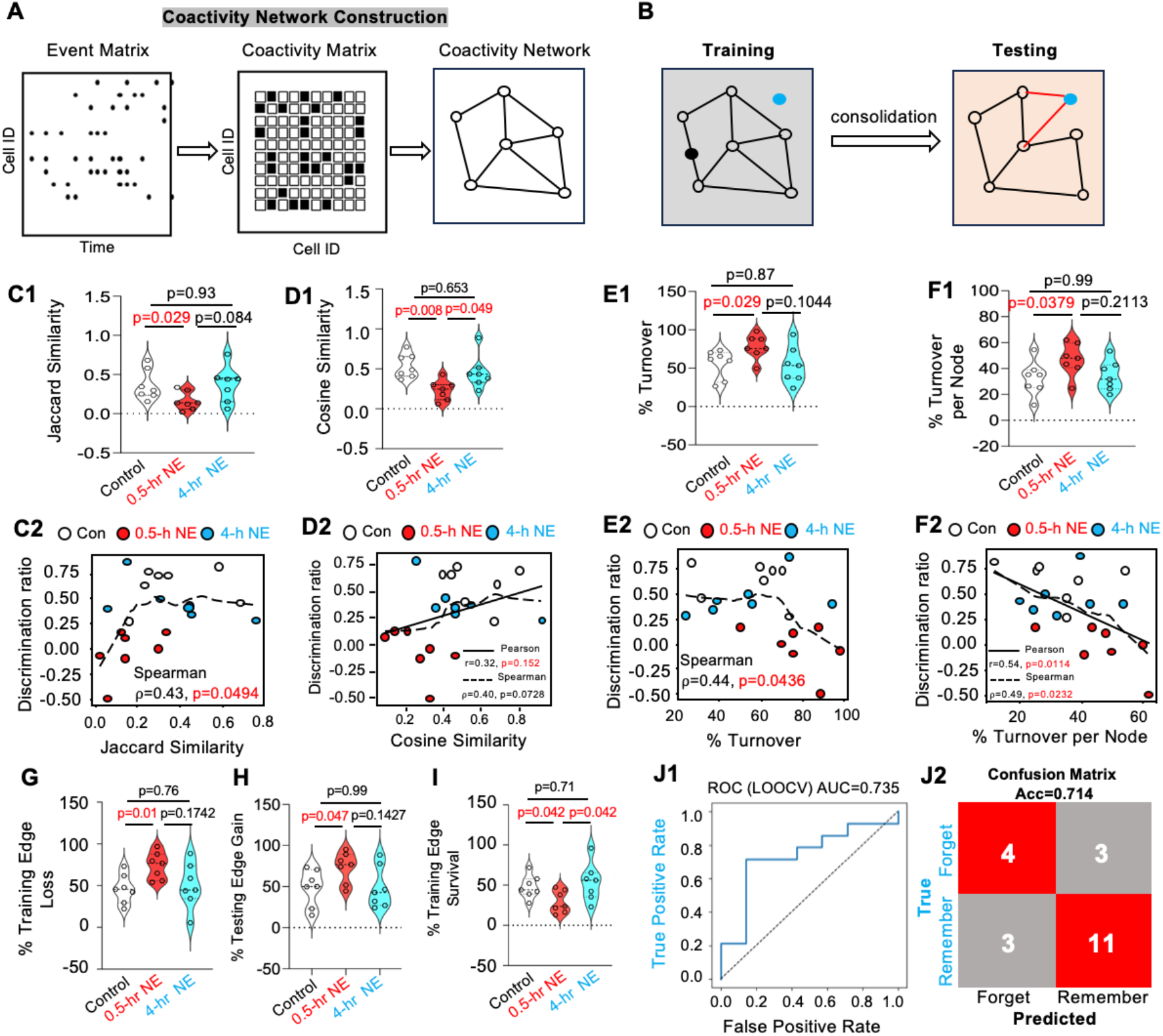
Temporally constrained RI disrupts the engram coactivity network. **(A)**. Schematic of coactivity network construction. (**B)**. Schematic of coactivity network manifested during training and testing. (**C1)**. Jaccard similarity [F (2,18)=3.365, p=0.0464]. (**C2)**. Correlation analysis between discrimination ratio and Jaccard similarity. (**D1)**. Cosine similarity [F (2,18)=6.413, p=0.0079]. (**D2)**. Correlation analysis between discrimination ratio and cosine similarity. (**E1)**. Network turnover rate [F (2,18)=3.595, p=0.0476]. (**E2)**. Correlation analysis between discrimination ratio and turnover rate. (**F1)**. Per-neuron edge turnover rate [F (2,18)=2.77, p=0.0894]. (**F2)**. Correlation analysis between discrimination ratio and % edge turnover per-neuron. (**G)**. Loss of training-specific edges during testing [F (2,18)=3.595, p=0.0486]. (**H)**. % Gain of edge during testing [F (2,18) =2.918, p=0.0799]. **i**. Survival of training-specific edges during testing [F (2,18)=3.616, p=0.0487]. (**J1)** and **(J2)**. Machine learning using coactivity network metrics predicts behavioral outcomes above chance. (**J1)**. ROC curve for leave-one-out cross-validated classification of Remember vs Forget animals using multivariate network remodeling features that include Jaccard similarity, turnover rate per neuron, training edge survival rate, training edge loss rate, and testing edge gain rate. The LOOCV logistic regression classifier yielded an AUC of 0.735 (**J1**) and an overall accuracy of 71.4% (Acc=0.714) (**J2**). Data presented as mean +/-SEM (n=7 for each group). Data were analyzed by one-way ANOVA followed by post hoc pairwise comparison.

Jaccard similarity (*16*) quantifies the fraction of coactivity edges shared between training and testing relative to all edges observed across both sessions. The control group showed a similarity score of 0.35 ± 0.074 (**Fig. 6C1**), reflecting a combination of engram drifting and reorganization along with the new network emerged during testing (*17*). Remarkably, the 0.5-hr NE group showed significantly decreased engram coactivity similarity (**Fig. 6C1**). In contrast, as a non-effective RI, 4-hr NE did not affect similarity between the coactivity networks during training and testing (**Fig. 6C1**). Further, Jaccard similarity of the coactivity network positively correlated with the memory discrimination ratio by Spearman analysis (**Fig. 6C2**).

To determine whether preservation of the weighted coactivity structure relates to memory retrieval, we quantified the cosine similarity between weighted coactivity networks of training and testing (**Fig. 6D**). The control and 4-hr NE groups exhibited higher cosine values compared to the 0.5-hr NE group (**Fig. 6D1**), consistent with greater preservation of training-related network structure in conditions that support memory retention. Across all animals, cosine similarity was marginally positive correlated with the discrimination ratio (**Fig. 6D2**).

The effective but partially reactivated engram coactivity network upon memory retrieval is, in principle, due to drifting and turnover. The turnover rate quantifies the fraction of coactivity edges gained or lost across the training and testing sessions. The control group’s turnover rate is 56 +/-6.9% (**Fig. 6E1**), reflecting reorganization of functional interactions across training and testing or drifting. The 0.5-hr NE group showed a significant increase in turnover rate relative to controls, whereas the 4-hr NE group was indistinguishable from the control group (**Fig. 6E1**). The turnover rate was negatively correlated with the memory discrimination ratio (**Fig. 6E2**).

For individual neuronal connectivity (i.e., the edges of each node in the coactivity network), we quantified the mean edge turnover index (Ti). The control group showed that 32.47 +/-5.01% of the connections per neuron were replaced during testing (**Fig. 6F1**). The 0.5-hr NE group showed a trend toward higher Ti (p = 0.064), while the 4-hr NE group was comparable to controls (p = 0.79) (**Fig. 6F1**). Pearson and Spearman analyses confirmed a negative correlation between Ti and the memory discrimination ratio (**Fig. 6F2**).

We investigated dynamic coactivity factors contributing to turnover. Analyses of training edge loss (**Fig. 6G**) and testing edge gain (i.e., new connections detected during testing) (**Fig. 6H**) revealed that 0.5-hour NE caused significant increases in both metrics (**Fig. 6G-H**), whereas non-effective RI (4-hr NE) did not influence connectivity dynamics (**Fig. 6G–H**).

While edge turnover analysis revealed substantial reorganization of the learning network, such global metrics do not distinguish between indiscriminate edge replacement and selective preservation of the engram network. We therefore quantified the survival rate of training-session edges to determine how learning-established functional connections are maintained during memory retrieval (**Fig. 6I**). In the control group, 47.64 ± 5.36% of training edges survived and were reactivated during memory retrieval. The 0.5-hr NE, but not the 4-hr NE, decreased training-edge survival rate (**Fig. 6I**).

We then asked whether these coactivity network parameters could predict memory outcome (binary classification: forgetting vs. remembering) using elastic net logistic regression. The model was trained on Jaccard similarity, turnover rate, turnover rate per neuron, training-edge survival rate, training-edge loss, and testing-edge gain. It predicted the classified memory state above chance (**Fig. 6J1**, AUC = 0.735; **Fig. 6J2**, accuracy = 0.714), demonstrating that multivariate network remodeling features carry predictive information about whether a memory will be retained or forgotten.

Together, these results demonstrate that effective but not non-effective RI causes a destabilization of the engram coactivity network, revealing a time-dependent influence of RI on the reorganization of functional neuronal assemblies.

### Location A sub-ensemble and network coactivity

Because the object at location A represents a shared spatial element during both training and the OLM test, we examined whether the location A–associated neuronal ensemble (**Fig. 7**) and its corresponding coactivity network (**Fig. 8**) are relevant to behavioral outcomes. Location A ensemble activity was defined as calcium activity detected while animals explored the object positioned at location A, whereas non–location A ensemble activity was defined as activity recorded when animals explored other spatial regions of the training arena.

**Fig. 7.**
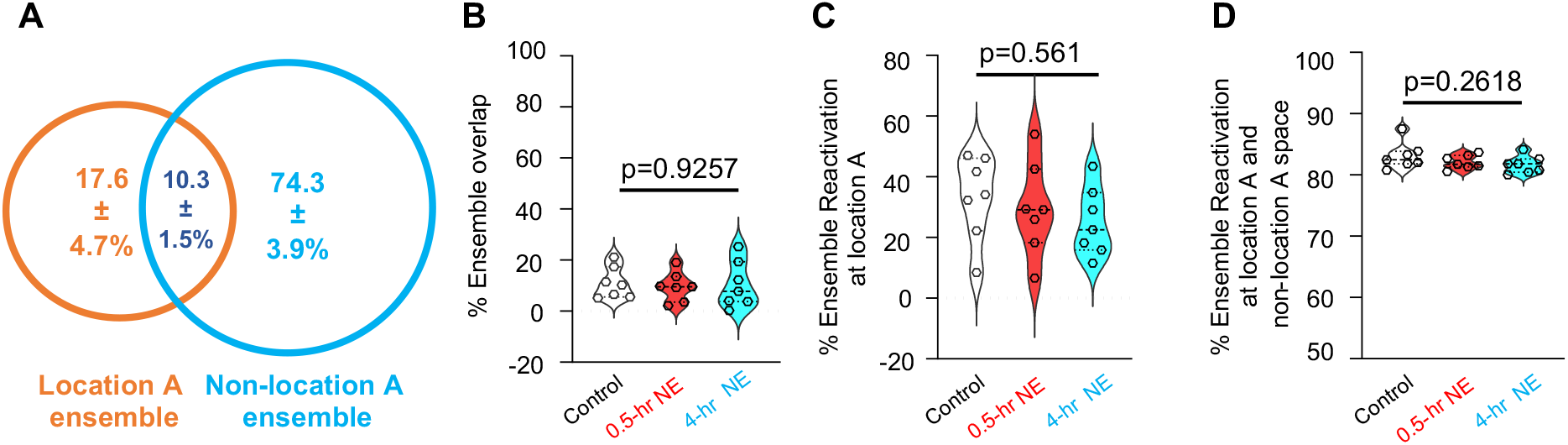
Location A ensemble activity during training and testing. Location A ensemble activity was detected by calcium imaging when animals explore the object at location A. Non-location A ensemble activity was detected when animals explore other spatial contents within the training chamber. (**A)**. Venn diagram showing the relative size and overlapping of location A and non-location A ensembles. (**B)**. Comparison of ensemble overlapping in distinct groups, including control mice and mice subjected to post-learning exposure to NE at 0.5 and 4 hr interval [(F (2,18)=0.0775, p=0.9257]. (**C)**. Location A ensemble reactivation when animals explore location A during OLM testing [(F (2,18)=0.6064, p=0.5561]. (**D)**. Location A ensemble reactivation when animals explore the whole spatial contents during OLM testing [F(2,18) =1.445, p=0.2618]. Data are mean +/-SEM and analyzed by one-way ANOVA (n=7 for each group).

**Fig. 8.**
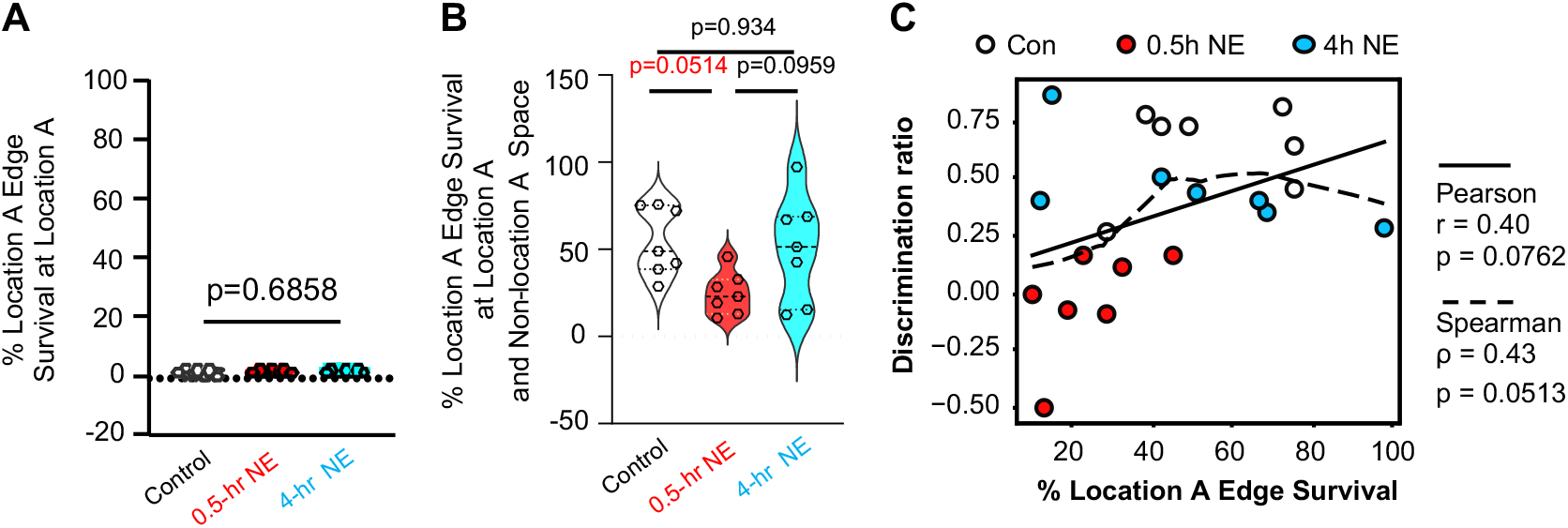
Location A-associated training edge survival. Within the ensemble network, connections (i.e., edges) manifested when animals explore location A during training were used as reference. Survival rate examines the degree of location A-associated edge reactivation during OLM testing (n=7 for each group). (**A)**. Location A-associated edge reactivation when animals explore location A during OLM testing [F (2,18)=0.3582, p=0.6858]. (**B)**. Location A-associated edge reactivation when animals explore the whole spatial contents during OLM testing [(F (2,18)=3.818, p=0.0415]. Data are mean +/-SEM and analyzed by one-way ANOVA followed by *post hoc* pairwise comparison (**A** and **B**). (**C)**. Pearson and Spearman analysis were used to identify correlation between memory discrimination ratio and location A-associated edge reactivation when animals explore the whole spatial contents during OLM testing.

We found that the location A sub-ensemble constituted 25.7 ± 3.5% of the total engram population. Ensemble overlap (10.3 ± 1.5%) was observed between neurons associated with location A exploration and those associated with non–location A spatial exploration (**Fig. 7A**). The degree of sub-ensemble overlap during training was comparable across the control, 0.5-hr NE, and 4-hr NE group (**Fig. 7B**).

During OLM testing, reactivation of the location A sub-ensemble was significantly lower when animals explored location A (i.e., the familiar location; 32.1 ± 5.2%, **Fig. 7C**) compared with when they explored other spatial regions (83.1 ± 0.85%, **Fig. 7D**). The extent of location A sub-ensemble reactivation did not differ significantly among control, 0.5-hr NE, and 4-hr NE group (**Fig. 7C** and **7D**).

At the network level, reactivation of location A–associated edges was minimal (1.5 ± 0.11%) when animals explored the familiar location during the OLM test (**Fig. 8A**). In contrast, when animals explored non–location A spatial regions, location A edge reactivation increased markedly to 54.5 ± 7.64% (**Fig. 8B**). This pattern is consistent with the behavioral principle underlying OLM performance, whereby successful memory recall is reflected by reduced exploration of the familiar location. Under these conditions, engram activity associated with the familiar location remains largely silent, whereas reactivation occurring in other spatial contexts may bias exploration away from the familiar site.

Notably, the 0.5-hr novelty exploration (NE) group exhibited significantly reduced location A edge reactivation compared with the control and 4-hr NE groups (**Fig. 8B**; *F*(2,18) = 3.818, *p* = 0.0415). Correlation analyses further revealed a marginal association between location A edge survival and memory strength, as measured by the location discrimination ratio (Pearson: *p* = 0.0762; Spearman: *p* = 0.0513; **Fig. 8C**). Together, these findings suggest that engram connectivity patterns, rather than ensemble membership alone, provide a functionally informative metric of memory-guided behavior.

### RI infiltrates the engram coactivity network

Although NE-specific neurons in both the 0.5-hr and 4-hr NE groups were reactivated alongside engram neurons during testing (**Fig. 4G**), only 0.5-hr NE affected memory retention (**Figs. 1A, 3** and **4C**) and overall coactivity network similarity and turnover (**Fig. 6C–H**). To investigate how NE-specific neurons functionally influence the coactivity network, we examined the topological structure of the engram network using network centrality metrics (*18, 19*), which quantify the roles of individual engram cells within the network and reflect a functional hierarchy (**Figs. 9A1** and **9B1**).

**Fig. 9.**
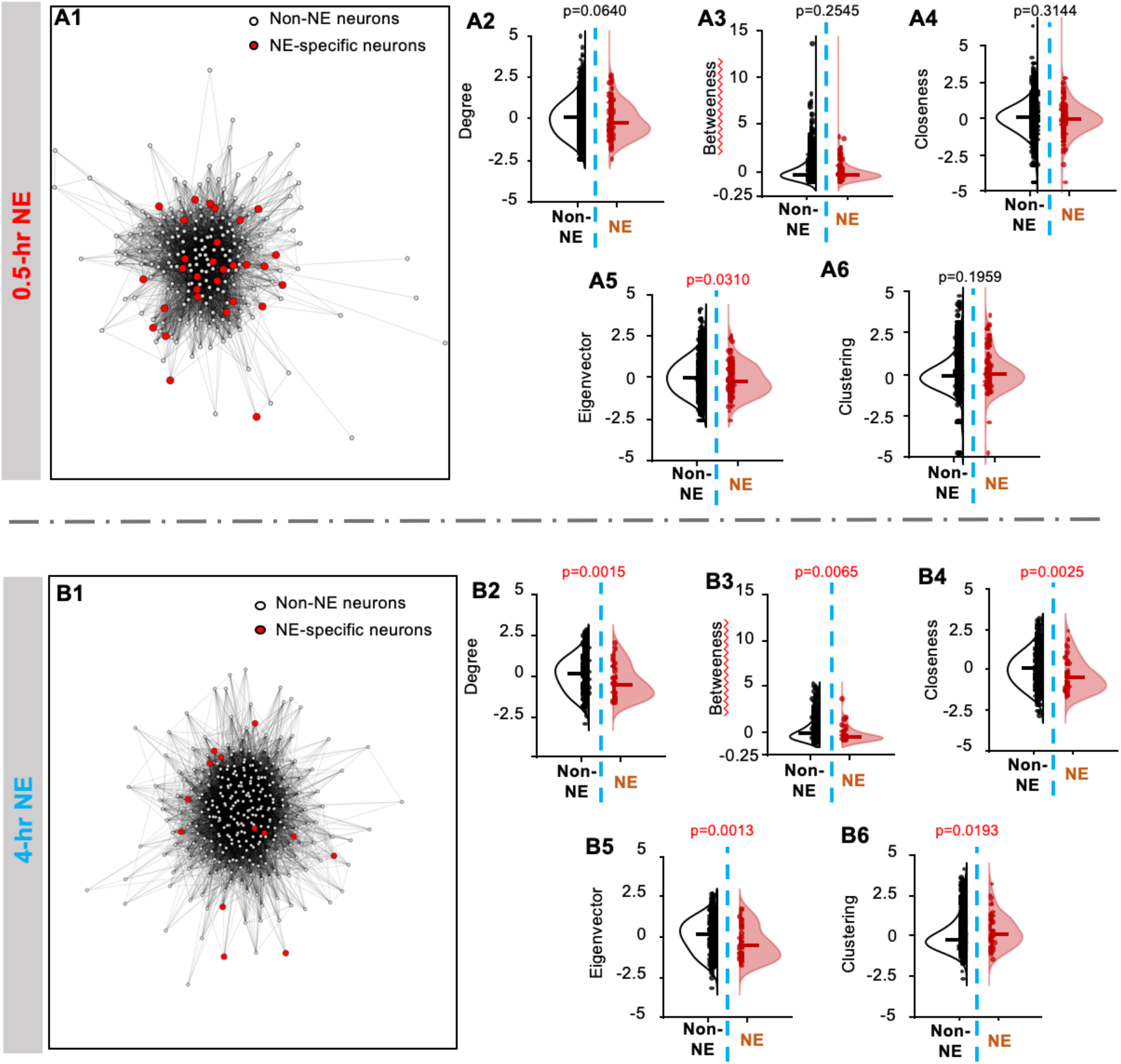
RI causes contamination of the core structure within the engram coactivity network. **(A1)**. Representative coactivity network manifested during testing in the 0.5-hr NE group. (**B1)**. Representative coactivity network manifested during testing in the 4-hr NE group. Both networks consist of NE-specific neurons and non-NE neurons as indicated. Comparison of the relative topological positions of the NE (n=149 for the 0.5-hr NE group; n=54 for the 4-hr NE group) and non-NE neurons (n=2823 for the 0.5-hr NE group; n=1473 for the 4-hr NE group) within the engram network was analyzed by z-scored degree (**A2** and **B2**), z-scored betweenness (**A3** and **B3**), z-scored closeness (**A4** and **B4**), z-scored eigenvector (**A5** and **B5**), and z-scored clustering (**A6** and **B6**). The Mann-Whitney test was used for statistical analysis.

Degree quantifies the number of edges per node, reflecting the overall connectivity of each engram cell (**Figs. 9A2** and **9B2**). Betweenness centrality quantifies how frequently a neuron lies on the shortest paths between other neurons, indexing its potential role in coordinating network-wide interactions (**Figs. 9A3** and **9B3**). Closeness captures the average proximity of a neuron to all others, reflecting its accessibility within the network (**Figs. 9A4** and **9B4**). Eigenvector centrality assigns higher scores to neurons connected to other highly connected neurons, identifying nodes embedded within influential network cores (**Figs. 9A5** and **9B5**). The clustering coefficient measures the degree to which a neuron’s neighbors are also interconnected, reflecting local network cohesiveness (**Figs. 9A6** and **9B6**).

Overall, the effective RI (i.e., the 0.5-hr NE) infiltrates the engram and contaminates the center core of the coactivity network; the non-effective RI (i.e., the 4-hr NE) infiltrates the engram and contaminates the peripheral structure of the coactivity network (**Fig. 9A1** and **9B1**). Within the coactivity network, the reactivated RI neurons (i.e., NE-specific neurons) in the 0.5-hr NE group intermingled with other engram cells (i.e., the non-NE neurons) and were indistinguishable from learning-activated engram cells (**Fig. 9A1** to **9A6**). In contrast, NE-specific neurons in the non-effective RI condition (4-hr NE) remained functionally distinct from other engram cells: they were less connected to the broader engram (**Fig. 9B2**), had longer connection paths to other engram neurons (**Fig. 9B3**), less accessible within the network (**Fig. 9B4**), preferentially associated with low-connectivity engram nodes in both the full and local coactivity networks (**Figs. 9B5–6**), and occupied peripheral network positions (**Fig. 9B1**). Thus, non-effective RI infiltrates the engram topologically but does not produce functional contamination of the coactivity network. These findings reveal a temporal dissociation in the topological properties of RI-activated neurons.

### Maturation of the engram coactivity network

Although the content of RI (i.e., novelty exploration) was identical across groups, the timing of RI produced distinct behavioral and network outcomes, implying a time-dependent vulnerability of the engram (*20*). We hypothesized that the engram undergoes progressive post-learning maturation, potentially reflecting the network aspects of consolidation process. We compared post-learning engram network reactivation at 0.5 hr (**Fig. 10A**) and 4 hr (**Fig. 10B**) after learning. From 0.5 to 4 hr post-learning, the network density increased (**Fig. 10C**) and the connection distance shortened (**Fig. 10D**), indicating a time-dependent enhancement of global network connectivity.

**Fig. 10.**
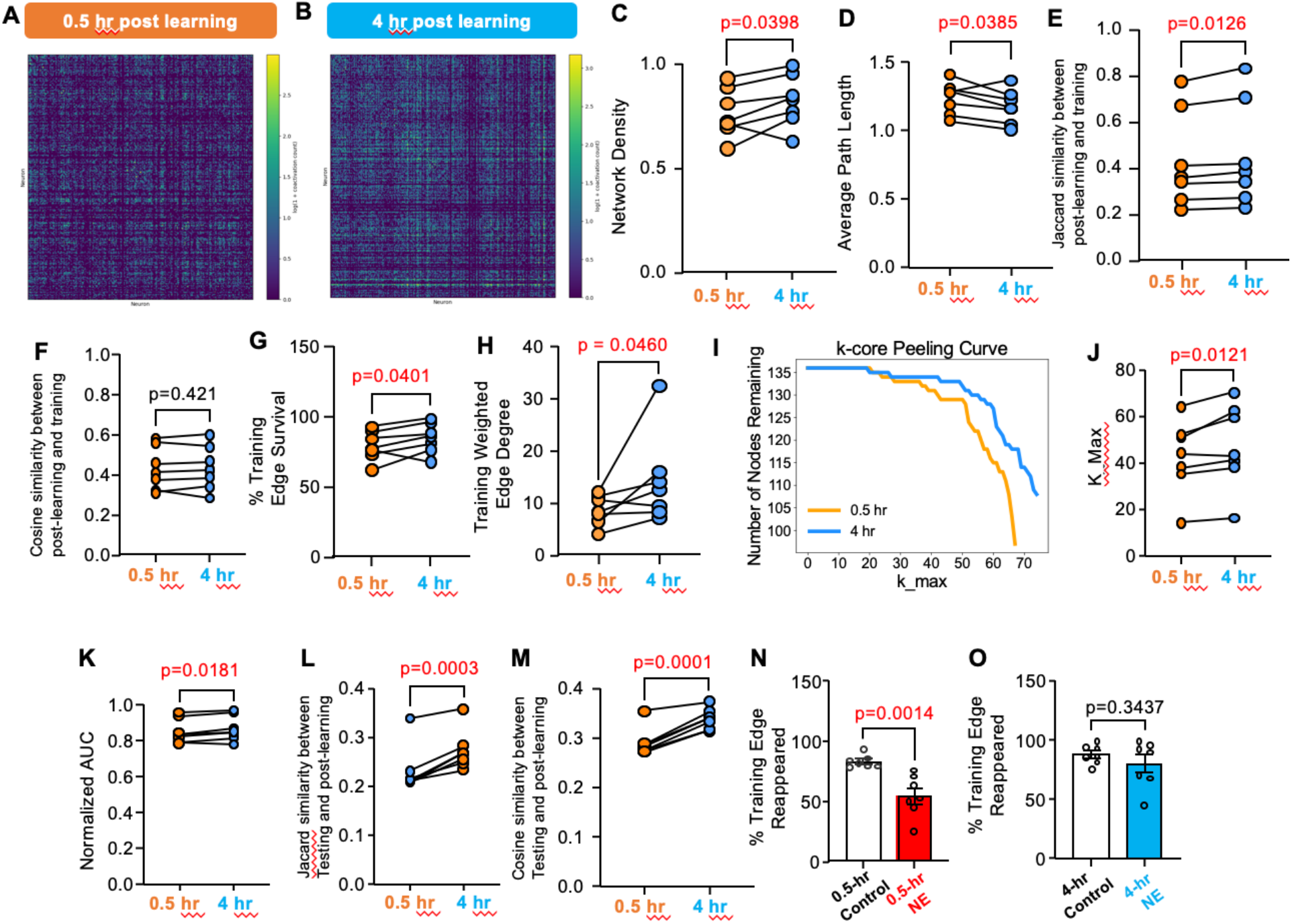
Progressive post-learning consolidation of the engram coactivity network. We examined the reactivated network in the same animal at 0.5 hr and 4 hr post-learning. **(A)** and (**B)**. Representative coactivity matrix detected 0.5 hr (**A**) and 4 hr (**B**) after learning. We compared coactivity network density (**C**), connection distance between nodes (**D**), Jaccard similarity (**E**), cosine similarity (**F**), survival of the connections established during learning (**g**), and coactivation frequency (**H**). Relevant to engram network structure, we compared the k-core peeling curve on training edges (**I**), k_max values associated with the reactivated network (**J**), and normalized AUC (area under the k-core peeling curve) (**K**) at 0.5 hr and 4 hr post-learning. Jaccard similarity (**L**) and cosine similarity (**M**) show that, comparing the engram network at 0.5 hr post-learning, the 4 hr post-learning network was preferentially reactivated during OLM testing. (**N)** Compared to controls, mice subjected to effective RI (i.e., NE at 0.5 hr after learning) show less learning engram reactivation during NE. (**O)**. Compared to controls, mice subjected to non-effective RI (i.e., NE at 4 hr after learning) show normal engram reactivation during NE. Data (n=7 for each group) were analyzed by paired two-tailed t-test (**C** to **H**; **J** to **M**) and unpaired t-test (**N** and **O**).

To determine whether the post-learning network reactivation is related to learning, we quantified both Jaccard similarity (**Fig. 10E**) and cosine similarity (**Fig. 10F**) of the networks manifested post-training (at 0.5 hr and 4 hr) versus training. Compared to the 0.5 hr post-training network, the 4 h post-training network exhibited higher similarity (**Fig. 10E**), although cosine similarity was not significantly altered (**Fig. 10F**).

To determine whether this global strengthening specifically consolidates training-related structure or merely reflects overall network densification, we isolated connections present during the training session. From 0.5 to 4 hr post-learning, training-defined edges showed significant increases in both edge presence (**Fig. 10G**) and coactivation frequency (**Fig. 10H**), suggesting progressive reinforcement of training-related functional connectivity. These findings indicate that training-derived edges become more deeply embedded and resistant to iterative node removal over time.

To directly assess whether the engram coactivity network undergoes structural maturation over time, we quantified network robustness using binary k-core decomposition (*21, 22*) at 0.5 and 4 hr after learning (**Fig. 10I**). Training-edge networks at 4 hr exhibited greater core depth and global robustness than at 0.5 hr. Specifically, the maximum core number (k_max) was significantly higher at 4 hr (**Fig. 10J**). Likewise, the normalized area under the k-core peeling curve (AUC), which captures distributed robustness across core levels, was also significantly increased at 4 hr (**Fig. 10K**).

The network maturation is functionally relevant. First, compared to the 0.5 hr post-learning network, the 4 hr post-learning network showed higher Jaccard (**Fig. 10L**) and cosine similarity (**Fig. 10M**) to the network manifested during memory retrieval. Second, upon network maturation, the learning-induced network was less disturbed. The 0.5-hr NE impaired the post-learning reactivation of the training-specific edges (**Fig. 10N**). The reactivation was normal at 4 hr post-learning with the concurrent NE exposure (**Fig. 10O**).

Taken together, our results suggest that post-learning engram activity reorganizes the training-derived coactivity scaffold into a structurally robust network architecture by 4 hours after learning, rendering the memory engram resistant to RI infiltration.

### Blocking RI infiltration rescues OLM

We used inhibitory DREADDs (i.e., hM4Di) to block neuronal activity within dHip during NE (**Fig. 11A** and **11B**). Strikingly, silencing post-learning neuronal activity in dHip using abolished both NE-induced neuronal activation (**Fig. 11C**) and NE-induced forgetting (**Fig. 11D** to **11F**). Mice expressing mCherry in the dHip and injected with CNO showed NE-induced forgetting (**Fig. 11D** to **11F**). Importantly, post-learning hM4Di activation did not affect novelty exploration per se; CNO-injected mice displayed normal locomotion in the novelty chamber (**Fig. 11G**) and normal object interaction during NE (**Fig. 11H**). Mice expressing either hM4Di or mCherry and injected with vehicle also showed NE-induced forgetting (**Fig. 12**). These results implicate a cellular mechanism in which local neuronal activation triggered by behavioral interference within a discrete brain region (i.e., the dHip) is necessary for activity-dependent forgetting of OLM.

**Fig. 11.**
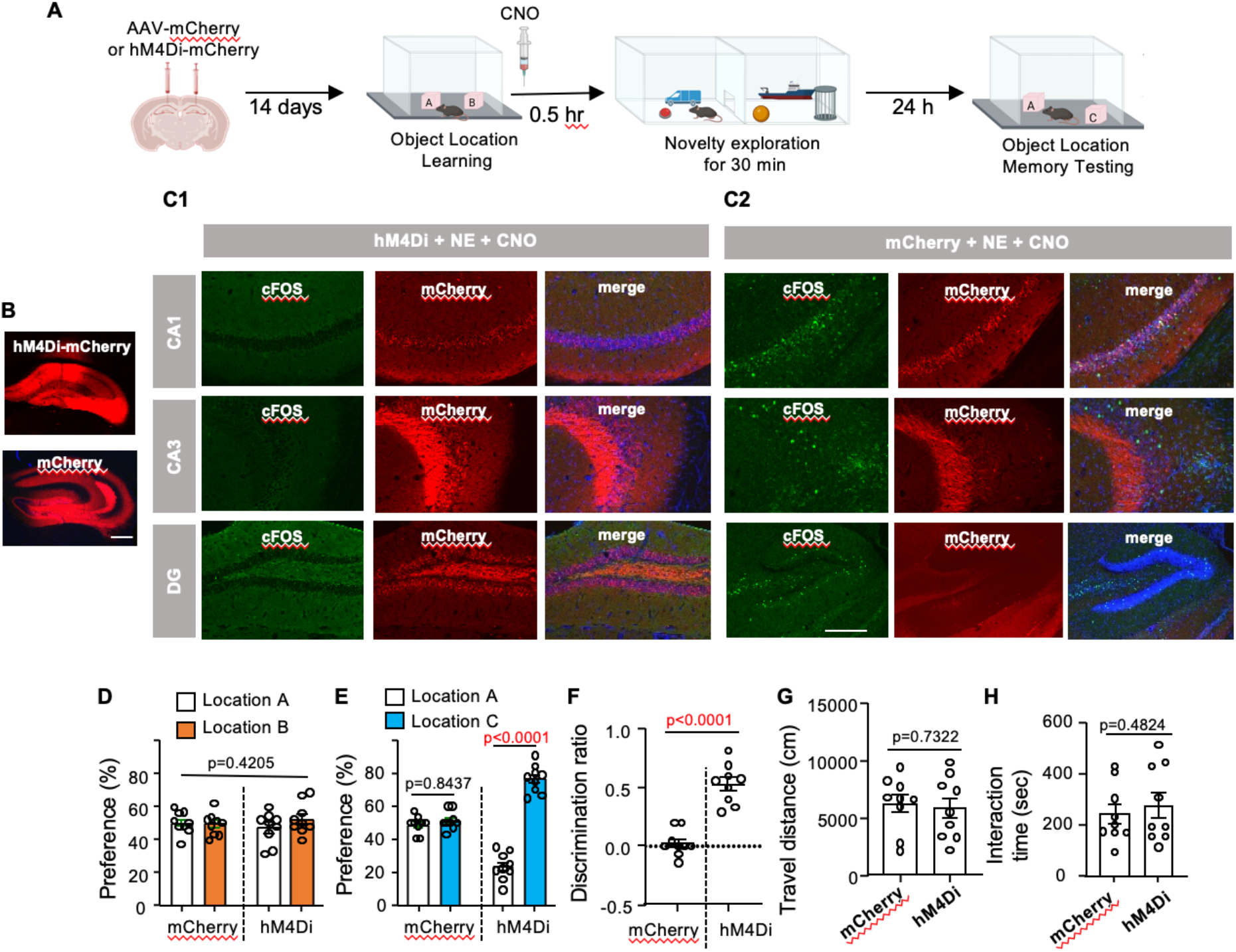
Inactivation of the dorsal hippocampus during post-learning NE rescues OLM. **(A)**. Procedures to express hM4Di or mCherry followed by learning, CNO injection paired with NE, and testing (n=9 for each group). (**B)**. Representative image of hM4Di-mCherry and mCherry expression in the dorsal hippocampus. Scale bar: 200 µm. (**C**). Mice expressing hM4Di-mCheery or mCherry were i.p. injected with CNO followed by a 30-min novelty exploration (NE). Brain sections were detected for mCherry along with immunostaining of cFos. While CNO administration blocked the NE-induced cFos upregulation in mice expressing h4MDi-mCherry (**C1**), there was no effect in mice expressing mCherry (**C2**). (**D)**. Object preference during learning [F (1,16)=0.6836 p=0.4205]. (**E)**. Object preference during testing [F (1,16)=55.65, p<0.0001]. (**F)**. Object discrimination during testing. (**G)** and **(H)**. Inactivation of the dorsal hippocampus during post-learning NE does not affect locomotion (**G**) and interaction with the novel objects (**H**). Data are presented as mean +/-SEM. Data were analyzed by unpaired t-test (**F, G**, and **H**) and two-way ANOVA (**D** and **E**) followed by post-hoc pairwise comparison.

**Fig. 12.**
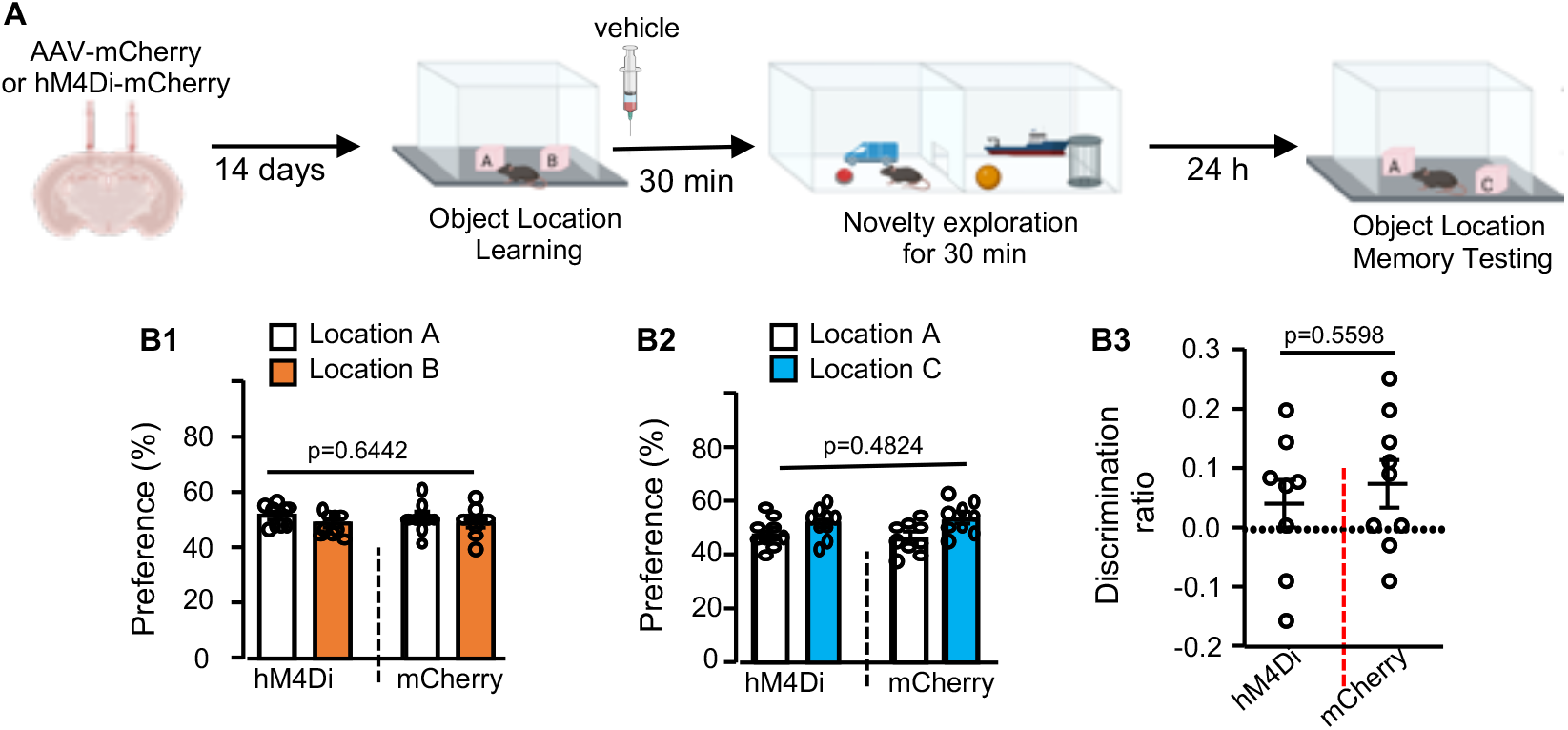
Vehicle administration in hM4Di-mCherry or mCherry expressing mice has no effect on NE-induced forgetting. **(A)**. Mice first received injection of AAV-hM4Di or mCherry into the dorsal hippocampus. 14 days after virus injection, mice were trained to learn 2 objects at locations A and B. The trained mice were then i.p. injected with vehicle followed by a 30-min NE. Mice were tested for OLM 24 hr later. Both groups (n=9 for the mCherry group; n=8 for the hM4Di group) showed normal and non-discriminative preference to object at location A and B during training [F (1.16) = 0.2216, p=0.6442] (**B1**). During OLM testing, both groups showed no preference to object at the new location C [F (1,16) =0.5173, p= 0.4824] (**B2**) and no discrimination ratio (**B3**). Data are presented as mean +/-SEM and analyzed by t-test (**B3**) and two-way ANOVA (**B1** and **B2**).

## DISCUSSION

### Neurobiology of forgetting

Despite decades of human cognitive research on RI-mediated forgetting, the neurobiology of the process is unknown. Classical interference theory proposed that forgetting arises primarily from the acquisition of competing associations rather than from passive trace decay (*23*). In our mouse model, post-learning dHIP activation is sufficient to cause forgetting. Further, silencing dHIP activity during NE abolishes forgetting without affecting exploratory behavior itself, providing strong evidence that RI-mediated forgetting reflects an active neuronal process, rather than passive degradation of the memory trace. This notion is consistent with studies showing that forgetting is mechanistically regulated (*1, 24, 25*).

One conceptually significant findings of this study is the precise temporal correspondence between the window of vulnerability to RI and the window of protein synthesis-dependent memory consolidation (*11, 12*). While some models proposed forgetting as a retrieval failure [(*26, 27*), but also see (*28*)], our findings offer an alternative conceptual framework. The forgetting mice show retrieval of reorganized and contaminated engram networks. Our findings are more consistent with a modification or consolidation disruption account. RI must occur during the consolidation window to impair memory, and effective interference is accompanied by measurable reorganization of engram coactivity networks along with infiltration of RI neurons rather than simply reduced retrieval probability. These results therefore support the interpretation that RI disrupts the stabilization of the underlying engram rather than merely preventing its retrieval.

### Functional metrics of the engram network

Recent advances in engram biology have demonstrated that memories are encoded in neuronal ensembles, which are functionally involved in memory storage and retrieval (*29, 30*). However, the current engram theory mainly focuses on engram membership (i.e., engram cells or ensembles) with little consideration of network coactivity (i.e., connections among engram cells). The finding that RI increases both training-edge loss and testing-edge gain suggests a two-sided disruption. The existing learning-related connections are weakened, and new interference-derived connections are incorporated. This is consistent with competitive synaptic plasticity models in which new synaptic weights overwrite or compete with previously formed weights (*31*). The present findings also support that connectivity stability, not just ensemble membership, is a determinant of memory persistence. The ability of a multivariate coactivity metrics model to predict binary memory outcome (AUC = 0.735) further validates this framework and raises the possibility that network topology could serve as a biomarker for memory vulnerability in physiological and pathological conditions.

### Structural hierarchy of the engram network

It has been suggested that the engram cells within the network are not functionally equal (*32*). A significant finding of the network analysis is the topological dissociation between effective and non-effective RI neurons within the engram coactivity network. NE-specific neurons activated during effective RI became structurally embedded in the core of the engram network, whereas NE-specific neurons activated during non-effective RI remained peripheral and topologically distinct. This “core infiltration” model provides a mechanistic explanation. Rather than the identity of the interfering experience, the degree to which the RI neurons are capable of integrating into the structural core of the engram determines whether forgetting occurs.

The finding that non-effective RI neurons infiltrate the engram peripherally, being less connected, more distant, and associated with low-connectivity nodes, suggests a gradient model of RI integration rather than a binary inclusion/exclusion model. This is consistent with computational models of associative memory networks in which the disruptive effect of new inputs scales with the degree to which those inputs overlap with the existing attractor state (*33*). In these models, memories are represented as stable attractor states within recurrent neural networks, and interference occurs when new activity patterns partially overlap with an established attractor, potentially destabilizing or modifying the stored representation (*34, 35*). Peripheral integration may therefore represent a form of partial interference that is insufficient to destabilize the core attractor but may still alter network dynamics at the margins.

### Engram maturation during the protein synthesis sensitivity window

The time-course of engram maturation, characterized by increasing network density, network similarity, connection dynamics, and k-core robustness between 0.5 and 4 hours post-learning, provides compelling evidence that consolidation operates not merely at the level of individual synapses but at the level of ensemble network architecture. This finding represents a conceptual extension of classical consolidation theory into the domain of functional network organization. Prior molecular and cellular studies of consolidation have focused predominantly on the stabilization of individual synapses through protein synthesis, intracellular signaling, and AMPA receptor trafficking (*36, 37*). The present data suggest that progressive integration and structural hardening of multi-neuron coactivity patterns represent an equally important aspect of consolidation.

The coincidence of network maturation with increasing resistance to both interference and protein synthesis inhibition suggests that structural consolidation of the ensemble and molecular stabilization of individual synapses are not independent processes, but rather interdependent components of a unified consolidation program. The engram-level maturation is broadly consistent with systems consolidation theory (*38*), which proposes that newly acquired memories initially depend on hippocampal circuitry before gradually reorganizing across distributed cortical networks over longer timescales. However, the present findings likely reflect an earlier intra-hippocampal phase of consolidation in which ensemble-level architecture is progressively stabilized.

Functionally, the consequences of engram infiltration appear to depend critically on task interval. Within the engram consolidation window, RI infiltration into the OLM engram leads to disruptive reorganization of the coactivity network. Time-dependent engram infiltration may also offer potential mechanisms underlying memory linking, in which two tasks in close temporal proximity show ensemble overlap, potentially allowing engram infiltration (*39*). This framework unifies interference and linking within a common competitive allocation model governed by network plasticity dynamics. Intriguingly, memory linking may also occur outside the original consolidation window through mechanisms involving reactivation of previously consolidated engrams. Recent evidence suggests that highly salient or traumatic experiences (e.g., repeated foot shocks) can trigger reactivation of existing memory traces, allowing new experiences to become associated with older, already consolidated engrams (*40*). These findings suggest that engram overlap can arise through at least two distinct mechanisms: a consolidation-window mechanism in which newly encoded ensembles compete or interfere during network maturation, and a reactivation-driven mechanism in which previously stabilized engrams are re-engaged and integrated with new experiences.

In summary, this work demonstrates that retroactive interference disrupts memory by infiltrating and reorganizing the structural core of a maturing hippocampal engram network. By unifying classical interference theory with modern engram biology and network neuroscience, these findings establish a mechanistic framework for understanding how experience reshapes memory during consolidation.

## MATERIALS AND METHODS

### Animals

All experiments were conducted with 2-to 4-month old C57BL/6 mice. Mice were group-housed (3-5 mice/cage) under a 12-h light/dark cycle and given free access to food and water at the Michigan State University animal facility. Behavioral experiments were performed during the light cycle. The Institutional Animal Care and Use Committee approved all procedures that follow the international guidelines for the use and care of laboratory animals.

### Stereotactic virus injection Viral vectors

2-month-old male mice were anesthetized with 80mg/kg ketamine (Dechra, 100mg/ml) and 6mg/kg xylazine (MWI Animal Health, USA). The head was fixed in a stereotactic apparatus (Kopf Instruments, USA). The skull was exposed, and a craniotomy was performed. AAV (0.5 μl per injection site) was injected into the dorsal hippocampus the following coordinates: anteroposterior (AP), -2.0 mm from bregma; mediolateral (ML), ±1.3 mm; dorsoventral (DV), -2.0 mm. All microinjections were carried out using a 5 µL microliter Syringe with a 34-gauge needle (Cat.84541, Hamilton, USA). Following each injection, the needle was left in place for 5 additional minutes to prevent back diffusion from the needle track and was then slowly withdrawn.

### Viral vectors

AAV9-hSyn-mCherry (Addgene, # 114472), AAV9-hSyn-hM4Di-mCherry (Addgene, # 50475), and AAV9-hSyn-jGCaMP7f-WPRE (Addgene, # 104488) were purchased from Addgene.

### Drug administration

Clozapine-N-Oxide (CNO, Cat. No. 4936, Tocris, Bristol, UK) was dissolved in saline containing 1% DMSO. For activation of hM4Di, 2mg/kg was administered (*42*). 1%DMSO in saline was used as vehicle control. All drugs were administered intraperitoneally at a volume of 10 ml/kg and in a randomized order to each subject.

### Behavior studies

Mice were pre-handled and habituated in the behavior testing room for 3 days. All the behavior tests were conducted during the light cycle of the day, from 9:30 am to 4:30 pm. Mice were transported to the training room and habituated for at least half an hour before any behavioral experiments.

The Object location memory task was conducted in an arena (16 × 16 × 16 inch) with spatial cues on each wall of the arena. Mice were first habituated in the training arena for 10 min per day for 3 days. During the 10-minute training, the animals were allowed to freely explore the chamber and interact with 2 objects at distinct locations (i.e., location A and B). The trained animals were tested 24 hours later. During the 10 min testing, animals were allowed to explore 2 objects with one at the old location (i.e., location A) and one at a novel location (i.e., Location C). The interaction time with object at each location during training and testing was recorded. Nose approaching the object within 1 cm of distance and/or direct touching the object with the nose was considered interaction. Preference was defined as time interacting with each object / the total interaction time with all objects. The discrimination index is calculated from the differential interaction time / total interaction time.

Mice subjected to pre-learning or post-learning novelty exploration were introduced to a novelty arena consisting of two chambers connected with a open passage and filled with 6 objects that were randomly positioned and differ from each other in shape and size. The total locomotion and interaction time with the objects were recorded during novelty exploration.

### Fluorescent microscopy

Mice were transcardially perfused with cold 4% paraformaldehyde (PFA) in PBS. The brain was extracted and fixed overnight in 4% PFA at 4 °C and cryoprotected in 30% sucrose in PBS. 30-micron brain sections were used to detect cFos (antibodies were from Cell Signaling Tech; Cat: #2250), hM4Di, mCherry, and GCaMP7f. The immunoactivity and fluorescent signal were detected by confocal microscopy.

### In vivo calcium imaging recording

Mice were injected with AAV9-hSyn-jGCaMP7f-WPRE into the right dorsal hippocampus. Two weeks later, mice were implanted with a gradient index (GRIN) lens (Inscopix, Mountain View, CA) above above the dorsal CA1 at -2.2 mm AP, 2.1 mm ML, and -1.35 mm DV with a 9-degree angle toward the medial direction. Mice with *GRIN* lens were single housed to prevent lens damage. A miniature microscope (Inscopix) was used to collect Ca^2+^ imaging videos in behaving mice. Videos were captured using nVista (Inscopix) at 20 Hz in a 720 × 549-pixel field (1.1 microns/pixel).

### End-to-end CNMF-E preprocessing and event extraction

Raw microendoscopic calcium imaging movies were preprocessed using end-to-end CNMF-E (IDEAS, Inscopix Data Exploration, Analysis, and Sharing) to extract neuronal activity events for downstream analysis. Briefly, movies from each animal were first concatenated (typically 3–4 sequential movies from the same session) into a single continuous time-series file to ensure consistent neuron identification and temporal modeling across the full recording. The concatenated movie was motion-corrected and then processed using CNMF-E to estimate neuronal spatial footprints (ROIs) and extract corresponding denoised fluorescence traces. Following CNMF-E, ROIs were manually inspected to exclude non-somatic components, duplicates, and low-quality signals. For longitudinal analyses, curated ROI sets from multiple sessions of the same animal were then aligned using cell registration (miniature correlation = 0.5) to identify matched neurons across sessions. After registration, neural events (Event Threshold Factor = 4, Event Smallest Decay Time = 0.2 second) were detected from extracted denoised activity traces using a consistent event-detection criterion and converted into a neuron-by-time neural event matrix for downstream analyses.

### Weighted Coactivity network construction (14)

The data were organized into a binary event matrix *E* ∈ ℝ^*T*×*N*^, where *T* denotes the number of time points and *N* the number of neurons. An entry *E*_*t,n*_ > 0 indicates that neuron *n*exhibited at least one detected calcium event at time point *t*; *E*_*t,n*_ = 0 indicates no event. To quantify functional coupling between neurons, we constructed a weighted, undirected coactivity network based on temporal co-occurrence of events. Coactivity was defined using a symmetric sliding window of 1 s (10 consecutive frames), advanced in steps of one frame (0.1 s) across the entire session. For each window position *w* (covering frames *t*to *t* + 9), we identified the subset of neurons that exhibited at least one event within that window. For every unordered neuron pair (*i, j*) (with *i* ≠ *j*) that were both active within the same window, the coactivity count for that pair was incremented by 1. This procedure was repeated across all window positions throughout the session. The result was a coactivity count matrix: *C* ∈ ℝ^*N* × *N*^, where each entry *C*_*ij*._ represents the total number of 1 s windows in which neurons *i* and *j* coactivated. The matrix *C* was interpreted as a weighted, undirected adjacency matrix, where edge weights correspond to coactivity counts (i.e., the frequency of temporal coactivation within the defined window). Thus, stronger functional coupling between two neurons is reflected by higher edge weight. The resulting graph for each session is defined as: *G* = (*V, E, W*), where: *V* is the set of active neurons, *E* is the set of undirected edges connecting coactive neuron pairs, *W* denotes the edge weight function *W*(*i,j*) = *C*_*ij*_. This procedure yields a session-specific weighted, undirected functional coactivity network suitable for subsequent graph-theoretic analyses.

### Isolation of training-specific edges

For each animal, training edges were defined as all neuron–neuron pairs that exhibited at least one coactivity event at any time during the entire training session. An edge (*i, j*) was classified as a training edge if it occurred at least once during the full training period [*E*_training_ = {(*i, j*) ∣ edge (*i, j*) occurred at any time during training }].

### Location A–specific edges

To isolate connectivity selectively associated with location A, we defined location A edges using temporal gating relative to location A occupancy. For each animal, behavioral timestamps corresponding to location A were identified. Edges that occurred within a 1 s window immediately preceding or during occupancy of location A were classified as location A–associated events. An edge (*i, j*) was defined as a location A–specific edge if it occurred at least once within this location A–aligned temporal window.

### Isolation of NE-specific neuron

We define NE-specific neurons as neurons that are preferentially active during the NE session but not engaged during Training sessions.

***Jaccard similarity and Turnover (16)*** were used to quantify how much the edge set changes between training (E1) and testing (E2): Jaccard 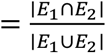. Accordingly, turnover was defined as 1 − Jaccard.

### Turnover rate per neuron

To quantify changes in network roles of individual neurons from training to testing, we computed turnover rate for each neuron: 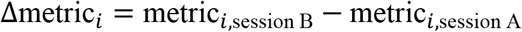.

### Training edge loss rate

The training edge loss rate was defined as the percentage of edges present during training that were absent during testing: Training edge loss rate 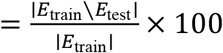.

### Testing edge gain rate

The testing edge gain rate was defined as the percentage of edges present during testing that were not present during training: Testing edge gain rate 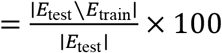.

### Training edge survival in testing

Training edge survival in testing was defined as the percentage of training edges that were retained in the testing network: Training edge survival 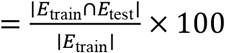.

### Classification model and cross validation strategy

To determine whether coactivity network properties predict behavioral memory outcome, we implemented supervised classification models to predict whether an animal remembered or forgot based on network-level features: Turnover rate, Jaccard similarity, turnover ate per neuron, training loss rate, testing gain rate and training edge survival rate to testing. Features were z scored across animals prior to model fitting to ensure comparable scaling. Memory outcome was defined behaviorally as control group and 4-hr NE group as remembering, 0.5-hr NE group as remembering. We used a logistic regression classifier to predict binary memory outcome (remember vs forget).The logistic model estimates: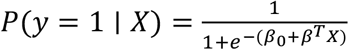 where: *y* = 1 indicates remembered, *X* is the feature vector, *β* are learned coefficients. Because of limited sample size, we used Leave-One-Out Cross-Validation (LOOCV): For each iteration: one animal was held out as test sample. Model was trained on remaining *n* − 1animals. Prediction probability was computed for the held-out animal. This process was repeated for all animals.

Performance Metrics. Model performance was evaluated using Accuracy: Accuracy 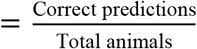. Confusion Matrix and ROC Curve and AUC. We computed the receiver operating characteristic (ROC) curve across prediction probabilities and calculated the area under the curve (AUC). AUC reflects the probability that a randomly chosen remembered animal receives a higher predicted probability than a forgotten animal.

### Centrality metric calculation (18, 19)

To compare NE and non-NE specific neuron’s role during testing, we computed node-level (per-neuron) metrics during testing coactivity network. From the constructed coactivity matrix *W*, where *W*_*i j*_ equals the number of windows in which neurons *i* and *j* were coactive (*W*_*ii*_ = 0).

1. Degree. Degree quantifies how many unique coactive partners a neuron has: *k*_*i*_ = Σ_*j* ≠ *i*_ *W*_*i j*_.
2. Betweenness centrality Betweenness measures how often neuron *i* lies on shortest paths between other neuron pairs: 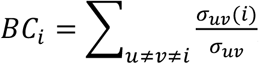, where *σ*_*uv*_ is the number of shortest paths between nodes *u* and *v*, and *σ*_*uv*_ (*i*) counts those paths that pass through *i*.
3. Closeness centrality Closeness captures how close neuron *i* is to all others in terms of shortest-path distance: 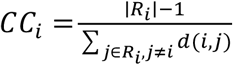where *R*_*i*_ is the set of nodes reachable from *i* and *d*(*i, j*)is the shortest-path distance.
4. Eigenvector centrality Eigenvector centrality assigns higher scores to neurons connected to other highly connected neurons: *E C*_;_ ∝ Σ_*j*_*W*_*ij*_ *EC*_*j*_, where *EC*_*j*_= centrality score of neuron *i, EC*_*j*_= centrality score of one of its neighbors.
5. Clustering coefficient Clustering coefficient quantifies the density of triangles around neuron *i*(i.e., whether its neighbors also coactivate with each other): 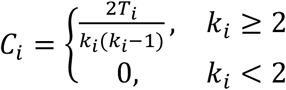 where *T*_*i*_ is the number of triangles that include node *i* in the graph, *k*_*i*_is the **number of neighbors** (coactive partners) of neuron *i*.

All node-level metrics were z-scored within each session to allow comparison across sessions and animals: 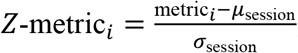, where *µ*_session_ and *σ*_session_ are the mean and standard deviation of that metric across all neurons in the given session.

### Global Network and training related edges Metrics

Network density.

Density was determined by Density 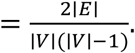, where E = number of edge (number of neuron pairs), V = number of nodes (number of neurons included in the network).

Average path length calculation. The average path length was computed as: 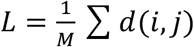 where: *d*(*i, j*)= shortest-path distance between neurons *i* and 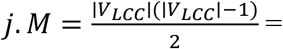total number of unordered node pairs in the largest connected component. Thus, *L* represents the mean minimum number of steps required to connect any two neurons in the network.

Weighed degree to training related edges. We computed restricted strength: 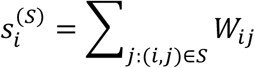, and normalized by session duration. *S* = training set of edges. The sum includes only edges belonging to *S*.

### k-core Decomposition and Peeling Curve (21, 22)

k-core decomposition was applied to 0.5 hr and 4 hr after learning binary coactivity network graph using an iterative peeling procedure. For a given integer k, the k-core was defined as the maximal subgraph in which all nodes have degree ≥ k. Starting from k = 0, nodes with degree < k were recursively removed until no further removals were possible. The number of remaining nodes was recorded for each value of k.

The k was not fixed a priori; instead, k-core decomposition was performed for all k ≥ 0 until the graph became empty, yielding a full k-core peeling curve. From this curve, we derived the following robustness metrics:

k_max defined as the largest k for which the k-core contained at least one node, reflecting the maximum depth of the core hierarchy.

Normalized AUC of the peeling curve. To obtain a scalar robustness measure, we computed the area under the peeling curve (AUC) using trapezoidal numerical integration and normalized it by the maximal possible area for that network:

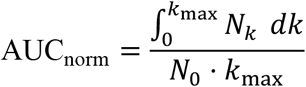

where *N*_0_is the number of nodes in the graph at *k* = 0(i.e., all shared neurons). AUC_norm_ranges from 0 to 1, with larger values indicating that a larger fraction of nodes persists across higher k thresholds. For each animal, k_max and normalized AUC were computed for both the 0.5-hr post-learning and 4-hr post learning sessions.

### Data collection, quantification and statistical analysis

The samples are biological replicates. Animals were used once and not repeated used for different experiments. The samples size is based on previous published results with OLM. The samples were randomly arranged and coded. When subjecting animals to NE, blinding is not possible. Blinding was practiced during OLM testing. For *in vivo* calcium imaging, sample exclusion is based on post-surgery complication, lack of or little GCaMP7f expression, and lens mis-alignment.

Statistical analyses were conducted using Prism (GraphPad) or Python (v3.12.6). Data distribution was assessed using the Shapiro-Wilk test. Parametric tests were used when data met normality assumptions; otherwise, non-parametric equivalents were applied. One-way and two-way ANOVA followed by post hoc pairwise comparison were used to analyze multiple groups with one or two factors. Unpaired and paired t-test were used to compare two groups. The p values were corrected for multiple comparisons where appropriate using standard correction methods. Specific statistic methods and sample size are provided in figure legends. Actual *F* and *P values* are reported in the main text or figure legends.

## Author contributions

H.W. conceived the study. L.A., M.Y., and H.W. designed the experiments. L.A. and M.Y. conducted experiments. L.A. analyzed the data. L.A. and H.W. wrote the manuscript.

